# A novel, complex-spike burst-dependent form of BCM-like metaplasticity regulates the induction of behavioral timescale synaptic plasticity

**DOI:** 10.64898/2026.07.17.739249

**Authors:** Thomas J. O’Dell

## Abstract

The induction of Hebbian LTP can produce a persistent, heterosynaptic suppression of LTP induction at other synapses. This form of metaplasticity, originally formalized in the Bienenstock, Cooper, and Munro (BCM) plasticity rule, generates competitive interactions between synapses and is thought to support sparse information encoding during memory formation. Importantly, although a non-Hebbian, burst-dependent form of synaptic plasticity known as behavioral timescale plasticity (BTSP) is essential for hippocampal memory encoding, little is known about the role of metaplasticity in BTSP. Thus, I examined whether the induction of BTSP at one set of synapses in the CA1 region of mouse hippocampal slices alters plasticity at other synapses. I find that the induction of BTSP by EPSP-evoked complex-spike (CS) bursts triggers a robust, but transient, heterosynaptic depression of excitatory synaptic transmission. This depression is induced by postsynaptic CS bursts, requires activation of L-type Ca^2+^ channels, and is mediated by activation of A1-type adenosine receptors. Notably, the heterosynaptic depression triggered by the induction of BTSP generates a CS burst-dependent form of BCM-like metaplasticity that transiently suppresses the induction of BTSP at other synapses. Together, these results provide experimental support for computational predictions that burst-dependent forms of plasticity are constrained by a distinct, burst-dependent form of BCM metaplasticity, and identify a mechanism that may contribute to sparse memory encoding during BTSP induction.

## Introduction

Although Hebbian forms of synaptic plasticity are thought to have a crucial role in learning and memory (Dringenberg, 2020), the computational power and behavioral relevance of Hebbian plasticity rules are limited by their unsupervised and correlative nature (i.e. changes in synaptic strength are triggered simply by the near coincidence of pre- and postsynaptic activity) (Gallistel and Matzel, 2013; Magee and Grienberger, 2020). The unsupervised nature of Hebbian plasticity is particularly problematic, as it allows new learning to erase previously encoded information by modifying the strength of synapses responsible for storing older memories, a phenomenon known as “catastrophic forgetting” (McClosky and Cohen, 1989; Ratcliff, 1990; French, 1999). Thus, in the absence of regulatory mechanisms that determine which synapses within a neural circuit are modified and when these changes occur, neural networks with Hebbian synapses will be highly susceptible to catastrophic forgetting (Zenke and Laborieux, 2024).

How can synapses use Hebbian plasticity to encode stable memories while avoiding catastrophic forgetting? Although multiple mechanisms are likely involved, different forms of metaplasticity may have an important role (Jedlicka et al., 2022). Metaplasticity encompasses a family of activity- and time-dependent mechanisms that regulate the ability of synapses to undergo Hebbian plasticity (Abraham, 2008). For example, although theta-frequency patterns of synaptic stimulation can erase LTP at recently potentiated synapses (i.e. induce depotentiation) (Fuji et al.,1991; O’Dell and Kandel, 1994), synapses become resistant to depotentiation within a few minutes after LTP induction (Staubli and Chun, 1996; Staubli and Scafidi, 1999). In addition, once potentiated, synapses enter a prolonged refractory period during which they are unable to undergo further potentiation (Frey et al., 1995; Moody et al., 1999; Kramar et al., 2012; Flores et al., 2025). Together, these homosynaptic forms of metaplasticity reduce the plasticity of potentiated synapses, thereby protecting them from undergoing catastrophic changes in synaptic strength during activity associated with the encoding of new memories. The induction of LTP can also trigger a heterosynaptic inhibition of LTP induction and facilitation of LTD induction at other synapses (Wang and Wagner, 1999; Abraham et al., 2001; Hulme et al., 2012, 2014). This heterosynaptic form of metaplasticity, first proposed in the Bienenstock, Cooper, and Munro (BCM) plasticity rule (Bienenstock et al., 1982), produces competitive synaptic interactions that can help prevent catastrophic forgetting by promoting sparse encoding of memories (Jedlicka et al., 2022).

Prior studies of metaplasticity have focused on its regulation of Hebbian forms of plasticity. Growing evidence indicates, however, that the encoding of both spatial (Magee, 2026) and non-spatial (Dorian et al., 2026) memories in the hippocampus relies on behavioral timescale synaptic plasticity (BTSP), a non-Hebbian form of synaptic plasticity induced by dendritic plateau potentials and postsynaptic complex-spike (CS) bursts (Bittner et al., 2017; Milstein et al., 2021; Madar et al., 2025). Although the mechanisms underlying BTSP and its functional properties are being extensively investigated, little is known about the role of metaplasticity in BTSP. Thus, it is unclear whether the same metaplastic processes that regulate Hebbian plasticity also influence BTSP, or whether BTSP is regulated by distinct forms of metaplasticity. Consistent with this latter possibility, computational studies using biologically realistic artificial neural networks suggest that burst-dependent forms of plasticity may be regulated by an unusual, burst-dependent variant of BCM metaplasticity (Payeur et al., 2021). Although this theoretical prediction has not yet been tested experimentally, the induction of BTSP at excitatory synapses on CA1 pyramidal cells is regulated by a burst-dependent form of synaptic competition induced by theta-frequency patterns of synaptic stimulation (O’Dell, 2022; 2025). Thus, in this study I examined the properties and mechanisms of theta-frequency stimulation-induced synaptic competition to explore its potential role as a novel, BCM-like form of metaplasticity.

## Results

The role of metaplasticity in the induction of BTSP was investigated using two stimulating electrodes to activate independent groups of Schaffer collateral fiber synapses (hereafter referred to as S1 and S2 synapses) in the CA1 region of hippocampal slices maintained in-vitro. A recording electrode placed in stratum radiatum between the stimulating electrodes was used to record field excitatory postsynaptic potentials (fEPSPs) and different patterns of theta-pulse stimulation (TPS) were used to elicit postsynaptic CS bursting and induce BTSP (O’Dell, 2022). As shown in figure 1, TPS-induced BTSP is strongly regulated by multiple forms of heterosynaptic plasticity (also see O’Dell, 2022; 2025). In these experiments, a 10-seconds-long TPS train was delivered to S2 synapses either alone (Fig. 1A) or following 10 or 30 seconds of TPS delivered to S1 synapses (Fig. 1B and C, respectively). In control experiments, where 10 seconds of TPS was delivered to S2 synapses alone, 25 ± 5% of the EPSPs evoked during S2 TPS elicited CS bursts (Supplemental Fig. S1A,B) and S2 fEPSPs potentiated to 131 ± 4% of baseline (n = 12) (Fig. 1A). In contrast, 97 ± 3% of the EPSPs evoked during S2 TPS elicited CS bursts when 10 seconds of S1 TPS was delivered before S2 TPS (Supplemental Fig. S1A,B) and S2 fEPSPs potentiated to 184 ± 4% of baseline (n = 11, *p* < 0.001 compared to S2 TPS alone; Fig. 1B). Thus although 10 seconds of TPS induces relatively modest levels of homosynaptic potentiation, this pattern of synaptic stimulation elicits a robust, heterosynaptic facilitation of both TPS-evoked CS bursting and BTSP induction. In contrast, increasing the duration of S1 TPS to 30 seconds induced a strong, heterosynaptic suppression of fEPSPs evoked during S2 TPS (Supplemental Fig. S1C,D) and inhibited the induction of BTSP at S2 synapses (45 min post-TPS S2 fEPSPs were just 105 ± 2% of baseline, n = 9, *p* < 0.001 compared to S2 TPS alone; Fig. 1C). Results from experiments where S1 TPS trains lasting 5-25 seconds were delivered before S2 TPS (Fig. 1D, Supplemental Fig. S2) indicate that although relatively brief trains of TPS (5-15 seconds duration) induce a heterosynaptic facilitation of BTSP induction, longer trains of TPS generate a potent, winner-take-all form of synaptic competition that suppresses the induction of BTSP at other synapses.

**Figure 1:**
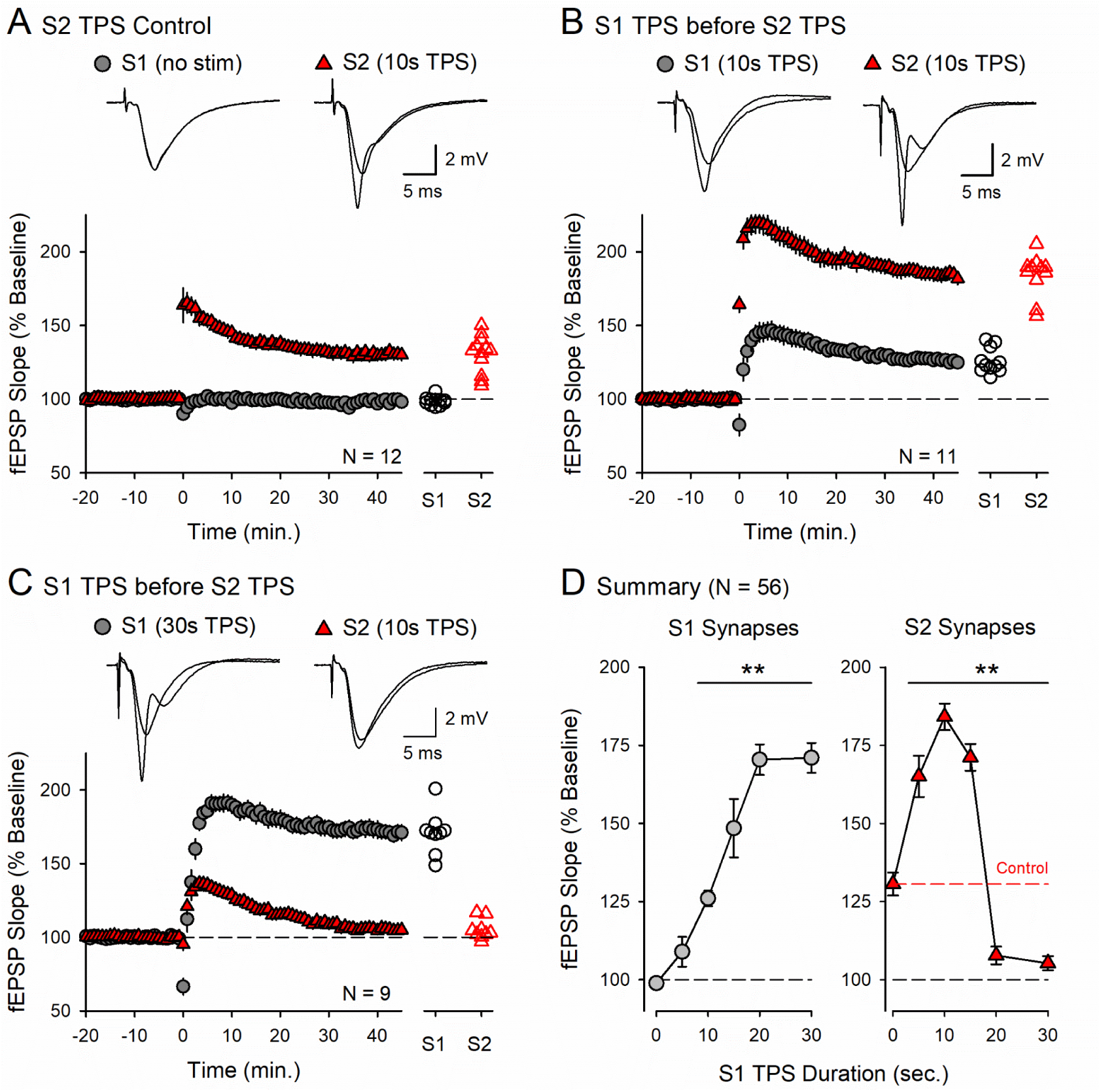
Multiple forms of heterosynaptic regulate the induction of BTSP. (***A***) Control experiments where 10 seconds of TPS was delivered to S2 synapses (at time = 0) without prior S1 TPS. 45 minutes-post TPS S2 fEPSPs were potentiated to 131 ± 4% of baseline and S1 fEPSPs were 99 ± 1% of baseline. (***B***) A brief train of S1 TPS (10 seconds duration) delivered before S2 TPS (10 seconds) facilitates BTSP induction at S2 synapses. S1 fEPSPs potentiated to 126 ± 3% of baseline and S2 fEPSPs were 184 ± 4% of baseline. (***C***) Increasing the duration of prior S1 TPS to 30 seconds inhibits TPS-induced BTSP at S2 synapses. S1 fEPSPs potentiated to 171 ± 5% of baseline while S2 fEPSPs were just 105 ± 2% of baseline. Scatter plots in A-C show S1 and S2 fEPSP slopes 45 minutes-post TPS from all experiments. Traces show superimposed fEPSPs recorded during baseline and 45 minutes-post TPS. (***D***) Summary showing results from all experiments where S2 TPS was delivered alone or after 5 – 30 seconds of S1 TPS. Changes in synaptic strength at S1 (left) and S2 synapses (right) were analyzed separately using one-way ANOVAs (\*\**p* < 0.001 compared to S2 TPS alone, S1 synapses: *F*_(5,50)_ = 54.975, *p* < 0.001; S2 synapses: *F*_(5,50)_ = 74.208, *p* < 0.001). The red line shows control levels of potentiation induced by 10 seconds of S2 TPS alone. Other results summarized in (D) are provided in Supplemental figure S2.

### A transient, BCM-like form of metaplasticity regulates the induction of BTSP

Is the heterosynaptic suppression of BTSP induced by longer trains of TPS a form of BCM metaplasticity? In the BCM plasticity rule, changes in synaptic strength are regulated in a cell-wide manner by a modification threshold, θ_M_, that determines the frequency of postsynaptic spiking required for the induction of LTP and LTD (Bienenstock et al., 1982; Copper and Bear, 2012). Thus, synapses that trigger frequencies of postsynaptic spiking greater than θ_M_ potentiate while synapses that elicit frequencies of postsynaptic spiking below θ_M_ depress. A key feature of the BCM plasticity rule is that θ_M_ is dynamically adjusted by levels of postsynaptic firing such that the threshold for LTP induction increases (and the induction of LTD is enhanced) after a bout of high frequency postsynaptic spiking. Importantly, this property of the BCM rule generates competitive interactions between synapses, i.e. the increase in postsynaptic activity evoked during the induction of LTP at one group of synapses triggers a heterosynaptic shift in θ_M_ that simultaneously suppresses LTP induction and facilitates the induction of LTD at other synapses. Thus, to determine whether TPS induces a BCM-like form of metaplasticity, I examined how the strength of S2 synapses is regulated by 100-pulse trains of 2-100 Hz stimulation delivered alone or after 30 seconds of TPS delivered to S1 synapses (Fig. 2, Supplemental Fig. S3). In control experiments, 2 Hz stimulation trains had no lasting effect on the strength of synaptic transmission (Supplemental Fig. S3A) while 3 Hz stimulation induced a modest potentiation (Fig. 2A). Higher frequencies of S2 stimulation induced more robust potentiation (Fig. 2B-D). Consistent with the notion that TPS induces a BCM-like form of metaplasticity, 30 seconds of S1 TPS delivered before S2 stimulation blocked the potentiation induced by 3 Hz stimulation (Fig. 2A) and significantly inhibited the potentiation induced by 5 Hz and 10 Hz stimulation (Fig. 2B,D). S1 TPS had no effect, however, on the potentiation of S2 synapses induced by 20 or 100 Hz stimulation protocols (Fig. 2C,D). Together, these results indicate that the induction of BTSP at S1 synapses produces a pronounced increase in the threshold for activity-dependent increases in synaptic strength at other synapses (Fig. 2D), as expected for a BCM form of metaplasticity.

**Figure 2:**
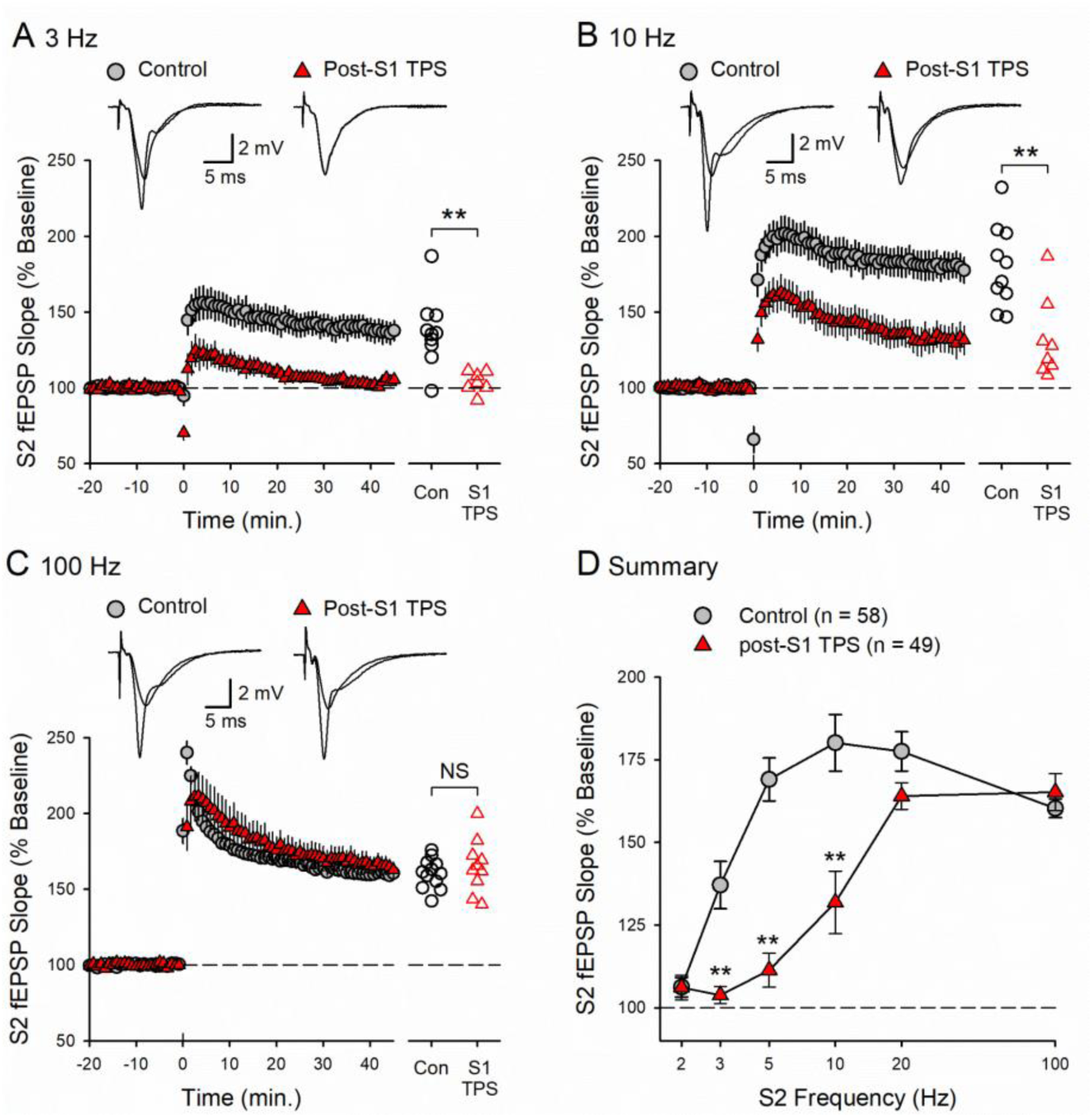
TPS induces a heterosynaptic, BCM-like shift in the threshold for potentiation at S2 synapses. (***A-C***) 100 pulses of S2 stimulation were delivered at 3 Hz (A), 10 Hz (B), or 100 Hz (C) either alone (control) or following a 30 seconds-long train of S1 TPS (stimulation trains delivered at time = 0). (***A***) S1 TPS significantly inhibited the induction of LTP induced by 3 Hz stimulation. S2 fEPSPs potentiated to 137 ± 7% of baseline in control experiments (n = 10) and were 104 ± 3% of baseline in experiments where S2 stimulation following S1 TPS (n = 7, *t*_(15)_ = 3.724, \*\**p* = 0.00203). (***B***) S1 TPS also inhibited the potentiation of S2 synapses induced by 10 Hz stimulation. S2 fEPSPs potentiated to 180 ± 9% of baseline in control experiments (n = 10) and were 131 ± 9% of baseline in experiments where 10 Hz stimulation followed S1 TPS (n = 8, *t*_(16)_ = 3.783, \*\**p* = 0.00163). (***C***) Prior S1 TPS had no effect on 100 Hz stimulation-induced LTP at S2 synapses. S2 fEPSPs potentiated to 160 ± 3% of baseline in control experiments (n = 12) and were 165 ± 6% of baseline in experiments where 100 Hz S2 stimulation followed S1 TPS (n = 10, *t*_(20)_ = 0.826, NS not significant, *p* = 0.418). Scatter plots in A-C show S2 fEPSP slopes 45 minutes post-100 pulse stimulation trains delivered to S2 synapses from all experiments and traces show superimposed S2 fEPSPs recorded during baseline and 45 minutes-post 100 pulse trains delivered to S2 synapses. (***D***) Summary for all experiments where 2 – 100 Hz trains of S2 stimulation (100 pulses) where delivered either alone (control) or after prior S1 TPS (duration = 30 seconds). A two-way ANOVA revealed a significant effect of S1 TPS (*F*_(1,95)_ = 51.656, *p* < 0.001), S2 stimulation frequency (*F*_(1,95)_ = 36.862, *p* < 0.001), and a significant interaction between S1 TPS and S2 stimulation frequency (*F*_(1,95)_ = 9.507, *p* < 0.001). S1 TPS significantly inhibited the potentiation induced by 3 Hz, 5 Hz, and 10 Hz S2 stimulation trains (\*\**p* < 0.001) but had no effect on changes in synaptic strength induced by 2 Hz (*p* = 0.991), 20 Hz (*p* = 0.105), or 100 Hz stimulation (*p* = 0.516) delivered to S2 synapses. Other results summarized in D are provided in Supplemental figure S3.

Notably, TPS trains that trigger a heterosynaptic suppression of BTSP also produced a robust heterosynaptic depression of excitatory synaptic transmission (Supplemental Fig. S1C,D). This is consistent with previous findings suggesting that heterosynaptic depression contributes to the TPS-induced inhibition of BTSP (O’Dell, 2022; 2025). However, TPS induces a transient form of heterosynaptic depression (O’Dell, 2025) and the strength of synaptic transmission at S2 synapses returns to baseline levels within ∼5 minutes following a 30 second train of S1 TPS (Supplemental Fig. S4A). This raised the possibility that the heterosynaptic suppression of BTSP induced by TPS might also be transient. To test this, I examined how BTSP induction at S2 synapses is affected when 30 seconds of TPS is delivered to S1 synapses at different intervals before S2 TPS. In control experiments, 100 pulses of TPS delivered to S2 synapses alone produced robust potentiation (S2 fEPSPs potentiated to 169 ± 6% of baseline, n = 8; Fig. 3A). When 30 seconds of S1 TPS was delivered immediately before S2 TPS, BTSP induction at S2 synapses was suppressed (45 min post-TPS S2 fEPSPs were 111 ± 5% of baseline, n = 8, *p* < 0.001 vs. control; Fig. 3B). In contrast, inserting a 5-minute interval between S1 and S2 TPS fully restored BTSP induction at S2 synapses (S2 fEPSPs potentiated to 176 ± 7% of baseline, n = 8, *p* = 0.43 vs. control; Fig. 3C,D). Thus, the heterosynaptic suppression of BTSP operates on the same short timescale as TPS-induced heterosynaptic depression, suggesting that this transient depression likely underlies the BCM-like suppression of BTSP induction.

**Figure 3:**
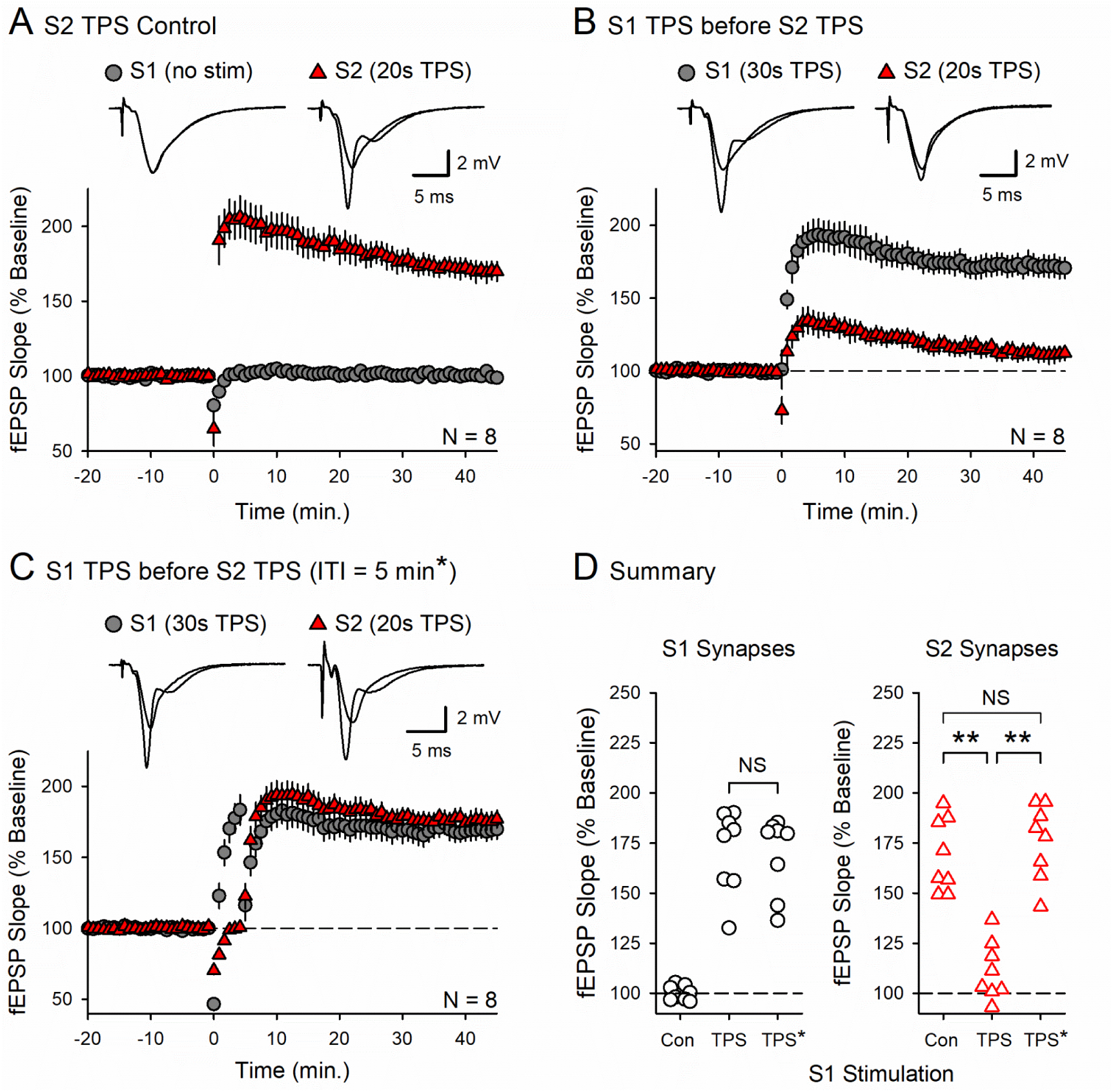
TPS induces a transient, heterosynaptic inhibition of BTSP. (***A***) Control experiments where 100 pulses of TPS was delivered (at time = 0) to S2 synapses. 45 minutes post-TPS S2 fEPSPs were potentiated to 169 ± 6% of baseline. (***B***) 30 seconds of S1 TPS delivered to S1 synapses before 100 pulses of S2 TPS potentiates S1 synapses and inhibits the induction of BTSP at S2 synapses. 45 minutes post-TPS S1 synapses were potentiated to 171 ± 7% of baseline and S2 synapses were just 111 ± 5% of baseline. (***C***) 30 seconds of prior S1 TPS does not inhibit BTSP induction at S2 synapses when TPS trains are delivered with a 5-minute inter-train interval (ITI). 45 minutes post-TPS S1 synapses were potentiated to 169 ± 7% of baseline and S2 synapses potentiated to 176 ± 7% of baseline. Traces in A-C show superimposed fEPSPs recorded during baseline and 45 minutes-post TPS. (***D***) S1 (left) and S2 (right) fEPSP slopes 45 minutes after TPS from all experiments in A-C. Changes in synaptic strength at S1 and S2 synapses were analyzed separately using one-way ANOVAs (\*\**p* < 0.001, NS not significant, S1 synapses: *F*_(2,21)_ = 49.027, *p* < 0.001; S2 synapses: *F*_(2,21)_ = 33.811 *p* < 0.001).

### TPS induces a CS burst-dependent form of heterosynaptic depression

To better define the potential role of CS bursting and heterosynaptic depression in the TPS-induced shift in θ_M_ that regulates BTSP, I next examined whether the heterosynaptic depression induced by TPS is a CS burst-dependent form of plasticity. Notably, EPSP-evoked CS bursting during TPS is highly activity-dependent and the number of EPSP-evoked CS bursts grows as the duration of TPS increases (O’Dell, 2022; Supplemental Fig. S4B). Thus, I first examined whether the magnitude of heterosynaptic depression at S2 synapses also grows as the duration of TPS delivered to S1 synapses increases (Supplemental Fig. S5). As shown in figure 4A and B, EPSPs-evoked during trains of S1 TPS lasting 15 or more seconds elicited CS bursts and a significant heterosynaptic depression at S2 synapses. In contrast, shorter trains of S1 TPS (5- and 10-seconds duration) produced little, if any, CS bursting and had no effect on transmission at S2 synapses (Fig. 4A,B). Consistent with the notion that TPS induces a CS burst-dependent form of heterosynaptic depression, the magnitude of heterosynaptic depression at S2 synapses was highly correlated with the probability of EPSP-evoked CS bursting during S1 TPS (*r* = −0.765, *p* = 4.426 x 10^-12^, n = 57; Fig. 4B).

**Figure 4:**
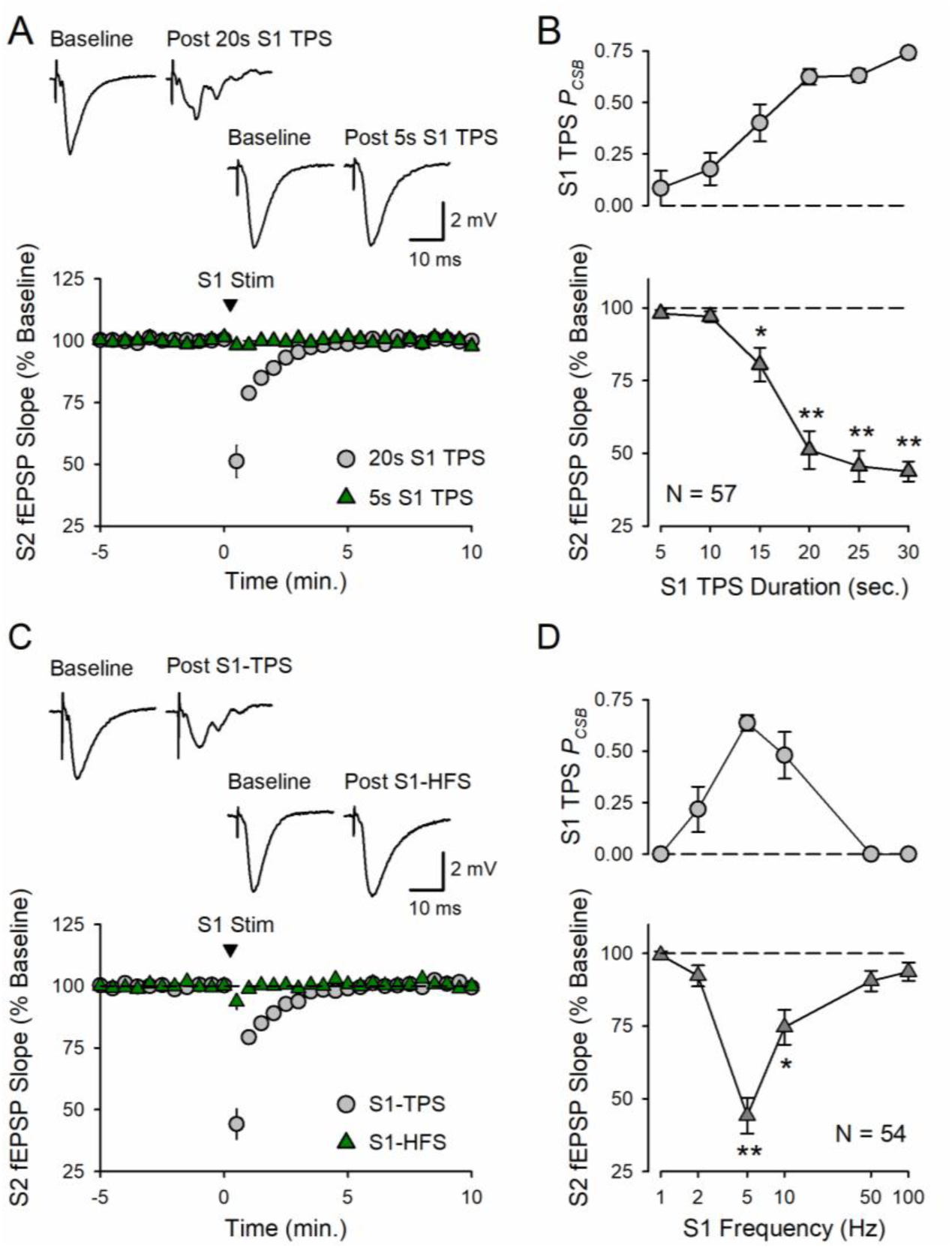
Heterosynaptic depression is selectively induced by patterns of synaptic activity that elicit postsynaptic CS bursting. (***A***) Examples of heterosynaptic depression induced by different durations of S1 TPS. Although 5 seconds of S1 TPS (delivered at time = 0) had little effect on transmission at S2 synapses (the first S2 fEPSPs elicited post-S1 TPS were 98 ± 1% of baseline, n = 9), a longer train of S1 TPS (20 seconds) induced a robust heterosynaptic depression at S2 synapses (the first S2 fEPSPs elicited post-S1 TPS were reduced to 51 ± 7% of baseline, n = 11). Traces show S2 fEPSPs recorded during baseline and after S1 TPS. (***B***) Summary showing the probability of EPSP-evoked CS bursting (*P_CSB_*) during S1 TPS (top) and the magnitude of heterosynaptic depression at S2 synapses (bottom) induced by different duration trains of S1 TPS. A One-way ANOVA revealed a significant effect of S1 TPS duration on the magnitude of heterosynaptic depression (*F*_(5,51)_ = 27.467, *p* < 0.001, \**p* < 0.05, \*\**p* < 0.001, compared to 5 seconds of S1 TPS). There was a statistically significant correlation between the *P_CSB_* during S1 TPS and the magnitude of heterosynaptic depression induced at S2 synapses (*r* = −0.765, *p* = 4.426 x 10^-12^). Additional results summarized in B are provided in Supplemental figure S5. (***C***) Examples of heterosynaptic depression induced by different frequencies of S1 stimulation. Although 100 pulses of S1 TPS (delivered at time = 0) inhibited transmission at S2 synapses (the first S2 fEPSPs evoked after S1 TPS were 44 ± 6% of baseline, n = 9), 100 pulses of high-frequency S1 stimulation (HFS, 100 Hz) had little effect on transmission at S2 synapses (the first S2 fEPSPs evoked after S1 HFS were 94 ± 3% of baseline, n = 10). Traces show S2 fEPSPs recorded during baseline and after HFS or TPS delivered to S1 synapses. (***D***) Summary showing the probability of EPSP-evoked CS bursting (*P_CSB_*) during S1 stimulation (top) and the magnitude of heterosynaptic depression at S2 synapses (bottom) induced by 100 pulse trains of S1 stimulation delivered at 1-100 Hz. A One-way ANOVA revealed a significant effect of S1 stimulation frequency on heterosynaptic depression (*F*_(5,48)_ = 21.548, *p* < 0.001, \**p* = 0.018, ** *p* < 0.001 compared to 1 Hz). The magnitude of heterosynaptic depression at S2 synapses was significantly correlated with the *P_CSB_* during S1 stimulation (*r* = −0.798, *p* = 4.976 x 10^-13^). Additional results summarized in D are provided in Supplemental figure S6.

Although the correlation between the probability of EPSP-evoked CS bursting during TPS and the magnitude of heterosynaptic depression (Fig. 4B) is intriguing, the magnitude of heterosynaptic depression observed in these experiments is also significantly correlated with the number of stimulation pulses delivered during S1 TPS (*r* = −0.81, *p* = 2.363 x 10^-14^). Thus, the correlation between CS burst probability and heterosynaptic depression, by itself, does not provide particularly compelling evidence for the notion that CS bursts trigger heterosynaptic depression. Thus, I next measured the magnitude of S2 heterosynaptic depression induced by trains of S1 stimulation containing the same number of stimulation pulses (100) delivered at 2 – 100 Hz (Supplemental Fig. S6). Importantly, the hypothesis that heterosynaptic depression is induced by postsynaptic CS bursts predicts that heterosynaptic depression at S2 synapses should only be induced by frequencies of S1 stimulation that elicit robust CS bursting. Consistent with this prediction, theta-frequency trains of S1 stimulation (5 and 10 Hz) elicited postsynaptic CS bursting and induced significant heterosynaptic depression (Fig. 4C,D). In contrast, little heterosynaptic depression was seen following lower frequencies of S1 stimulation (1 and 2 Hz) that elicited few or no CS bursts (Fig. 4D). Higher-frequency trains of S1 stimulation (50 and 100 Hz), which failed to elicit postsynaptic CS bursts, also had little effect on synaptic transmission at S2 synapses (Fig. 4C,D). Together, these results indicate that CS bursts provide a unique postsynaptic signal for triggering heterosynaptic depression. Consistent with this notion, there was again a significant correlation between the magnitude of heterosynaptic depression at S2 synapses and the probability of EPSP-evoked CS bursting during S1 stimulation in these experiments (*r* = −0.798, *p* = 4.976 x 10^-13^, n = 54; Fig. 4D).

As a more direct test of the role of CS bursts in heterosynaptic depression I next investigated whether enhancing EPSP-evoked CS bursting during TPS enhances the magnitude of TPS-induced heterosynaptic depression and, conversely, whether suppressing EPSP-evoked CS bursting inhibits the induction of heterosynaptic depression. Notably, dendritic plateau potentials and CS bursting are strongly suppressed by somatostatin-positive inhibitory neurons that target the apical dendrites of CA1 pyramidal cells (Lovett-Barron et al., 2012). Activation of mu-opioid receptors (μORs) suppresses the dendritic inhibition produced by these interneurons (Caccavano et al., 2025) and μOR agonists promote EPSP-evoked burst firing in CA1 pyramidal cells (Nicoll et al., 1980). Thus, I first examined whether facilitating EPSP-evoked CS bursting with the μOR agonist DAMGO enhances TPS-induced heterosynaptic depression.

In these experiments, a 10-minute bath application of DAMGO (1 µM) increased the amplitude of fEPSPs but had no effect on fEPSP slopes (Fig. 5A, Supplemental Fig. S7), suggesting that μOR activation suppresses inhibitory synaptic transmission in the CA1 region without altering excitatory synaptic transmission. DAMGO also enhanced EPSP-evoked CS bursting during a 20-seconds-long train of S1 TPS and significantly enhanced the heterosynaptic depression of S2 synapses induced by S1 TPS (the first S2 fEPSPs evoked post-S1 TPS in the presence of DAMGO were reduced to just 27 ± 2% of baseline compared to 53 ± 4% of baseline in interleaved control experiments) (Fig. 5A). In contrast, the heterosynaptic depression induced by 20 seconds of S1 TPS was abolished in experiments where a low concentration (200 nM) of the Na^+^ channel blocker tetrodotoxin (TTX) was bath applied for 10 minutes before S1 TPS to suppress EPSP-evoked CS bursting (the first S2 fEPSPs evoked post-S1 TPS in the presence of TTX were 98 ± 2% of baseline vs. 49 ± 5% of baseline in interleaved control experiments) (Fig. 5B). Thus, consistent with the notion that TPS induces a CS-dependent form of heterosynaptic depression, the magnitude of heterosynaptic depression is not only significantly correlated with the probability of EPSP-evoked CS bursting during different patterns of synaptic activity (Fig. 4) but is also regulated by pharmacological manipulations that alter dendritic excitability and EPSP-evoked CS bursting in CA1 pyramidal cells (Fig. 5).

**Figure 5:**
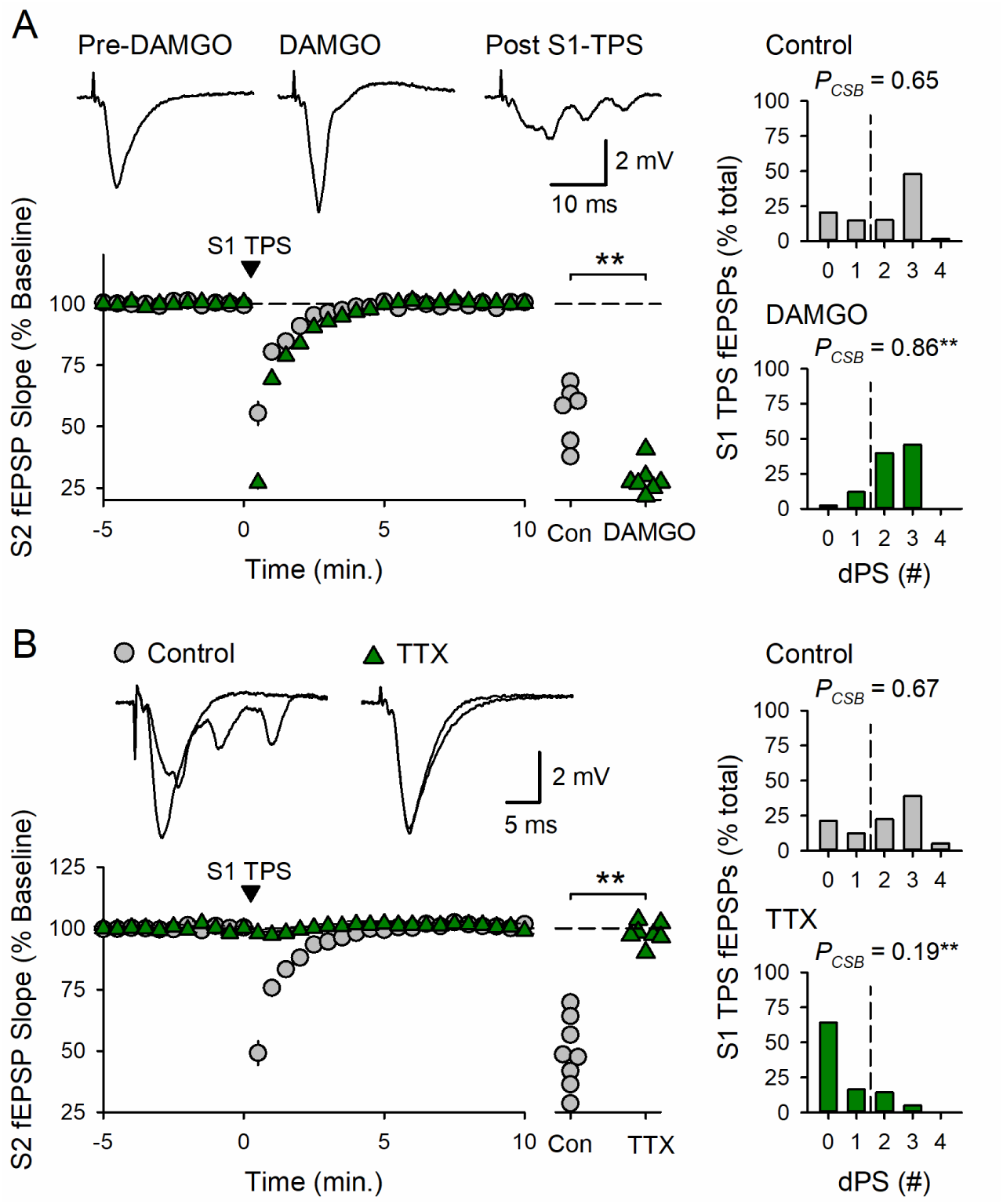
TPS induces a CS burst-dependent form of heterosynaptic depression. (***A***) The µOR agonist DAMGO (1 µM) increases TPS-induced CS bursting and enhances heterosynaptic depression. S2 fEPSPs were reduced to 53 ± 4% of baseline following 20 seconds of S1 TPS (delivered at time = 0) in control experiments (n = 8) and reduced to 27 ± 2% of baseline when S1 TPS was delivered at the end of a 10-minute bath application of DAMGO (n = 8). Scatter plots show the slopes of the first S2 fEPSPs evoked after S1 TPS from all experiments (*t*_(14)_ = 5.175, \*\**p* = 1.41×10^-4^). Histograms show the number of fEPSPs eliciting 0-4 dendritic population spikes (dPS) during S1 TPS from all experiments. DAMGO significantly enhanced EPSP-evoked CS bursting during S1 TPS (*t*_(14)_ = 5.862, \*\**p* = 2.52×10^-4^). Traces show example S2 fEPSPs evoked before DAMGO application, at the end of a 10-minute bath application of DAMGO, and the first S2 fEPSP evoked after S1 TPS in the presence of DAMGO. (***B***) Same experimental approach as in A but with results from experiments where S1 TPS was delivered at the end of a 10-minute bath application of TTX (200 nM). S2 fEPSPs were reduced to 49 ± 5% of baseline in control experiments (n = 8) and were 98 ± 2% of baseline following S1 TPS in the presence of TTX (n = 7, *t*_(13)_ = 8.887, \*\**p* = 6.95×10^-7^). TTX significantly inhibited EPSP-evoked CS bursting during S1 TPS (*t*_(13)_ = 5.852, \*\**p* = 5.66×10^-5^). Traces show superimposed S2 fEPSPs recorded during baseline and after S1 TPS.

### Activation of A1-type adenosine receptors mediates the BCM-like metaplastic suppression of BTSP

How might a CS burst-dependent form of heterosynaptic depression generate a BCM-like form of metaplasticity? One well established activity-dependent form of heterosynaptic depression at excitatory synapses onto CA1 pyramidal cells is due to an inhibition of glutamate release produced by activation of presynaptic A1-type adenosine receptors (Grover and Teyler, 1993; Mitchell et al., 1993; Wu and Saggau, 1994). Moreover, repetitive postsynaptic spiking triggers an L-type calcium channel-dependent increase in intracellular adenosine levels that drives the release of adenosine from CA1 pyramidal cells dendrites via equilibrative nucleoside transporters (Lovatt et al., 2012; Wu et al., 2023; Wei et al., 2025). Thus, the TPS-induced shift in θ_M_ and suppression of BTPS induction may be mediated by an adenosine receptor-dependent form of heterosynaptic depression that, by inhibiting glutamate release, prevents the strong activation of postsynaptic NMDA receptors needed for the induction of BTSP. Consistent with this potential mechanism, the inhibition of S2 synapses elicited by 20 seconds of S1 TPS was significantly attenuated in slices bathed in ACSF containing the L-type channel blocker nimodipine (Fig. 6A). Moreover, TPS-induced heterosynaptic depression was almost completely blocked in slices bathed in ACSF containing the A1 receptor antagonist DPCPX (Fig. 6B).

**Figure 6:**
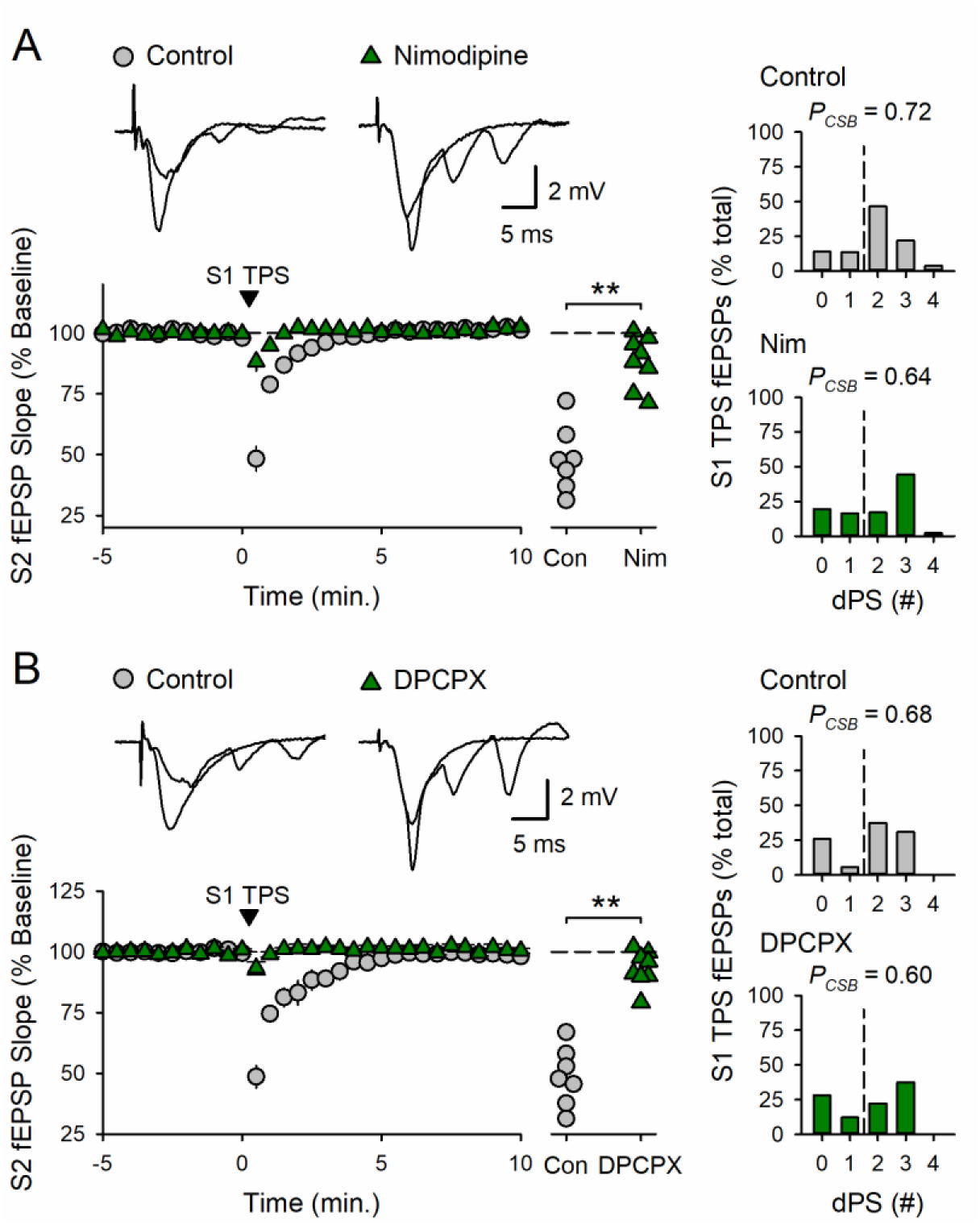
TPS induces an A1 receptor-dependent form of heterosynaptic depression. (***A***) The L-type Ca^2+^ channel blocker nimodipine inhibits TPS-induced heterosynaptic depression. In control experiments, S2 fEPSPs were initially depressed to 48 ± 5% of baseline (n = 7) after a 20-seconds-long train of S1 TPS (delivered at time = 0). In contrast, the first S2 fEPSPs evoked post S1 TPS were 88 ± 4% of baseline in experiments where slices were continuously bathed in ACSF containing 10 μM nimodipine (n = 8). (***B***) The A1-type adenosine receptor antagonist DPCPX inhibits TPS-induced heterosynaptic depression. S2 fEPSPs were initially depressed to 49 ± 5% of baseline following 20 seconds of S1 TPS (delivered at time = 0) in control experiments (n = 7) and were 93 ± 3% of baseline after S1 TPS in slices continuously bathed in ACSF containing 400 nM DPCPX (n = 8). Scatter plots in A and B show the slopes of the first S2 fEPSPs evoked following S1 TPS in all experiments. Both nimodipine (\*\**t*_(13)_ = 6.405, *p* = 2.33×10^-5^) and DPCXP (\*\**t*_(13)_ = 8.823, *p* = 7.52×10^-7^) significantly inhibited TPS-induced heterosynaptic depression. Histograms in A and B show number of EPSPs evoking 0 – 4 dendritic populations spikes (dPS) during S1 TPS in all experiments. Nimodipine (Nim) and DPCPX had no effect on the probability of EPSP-evoked CS bursting (*P_CSB_*) during S1 TPS (*t*_(13)_ = 1.384, *p* = 0.19 and *t*_(13)_ = 0.986, *p* = 0.342, respectively). Traces in A and B show superimposed S2 fEPSPs recorded during baseline and the first S2 fEPSP evoked post S1 TPS.

To determine whether TPS-induced heterosynaptic depression underlies the BCM-like metaplastic regulation of BTSP I next examined the effects of DPCPX on the induction of BTSP when 30 seconds of S1 TPS was delivered before 10 seconds of S2 TPS. Consistent with results shown in figure 1C, 30 seconds of S1 TPS induced BTSP at S1 synapses (fEPSPs potentiated to 165 ± 6% of baseline, n = 10) and suppressed the induction of BTSP at S2 synapses (45 minutes post-TPS, S2 fEPSPs were just 107 ± 6% of baseline) (Fig. 7A). In contrast, when this pattern of TPS was delivered in the presence of DPCPX, S1 synapses potentiated to 180 ± 6% of baseline and S2 synapses potentiated to 199 ± 6% of baseline (n = 9) (Fig. 7B). Thus, although DPCPX had no effect on the induction of BTSP at S1 synapses, blocking A1 receptors abolished the metaplastic suppression of BTP induction at S2 synapses (Fig. 7C) and revealed a strong heterosynaptic facilitation of BTSP normally only seen with shorter trains of S1 TPS (Fig. 1). DPCPX did not, however, facilitate the induction of BTSP when 10 seconds of TPS was delivered to S2 synapses without prior S1 TPS (Fig. 7D,E). This indicates that blocking A1 receptors does not simply enhance BTSP induction but instead selectively disrupts the BCM-like suppression of BTSP induced by 30 seconds of TPS.

**Figure 7:**
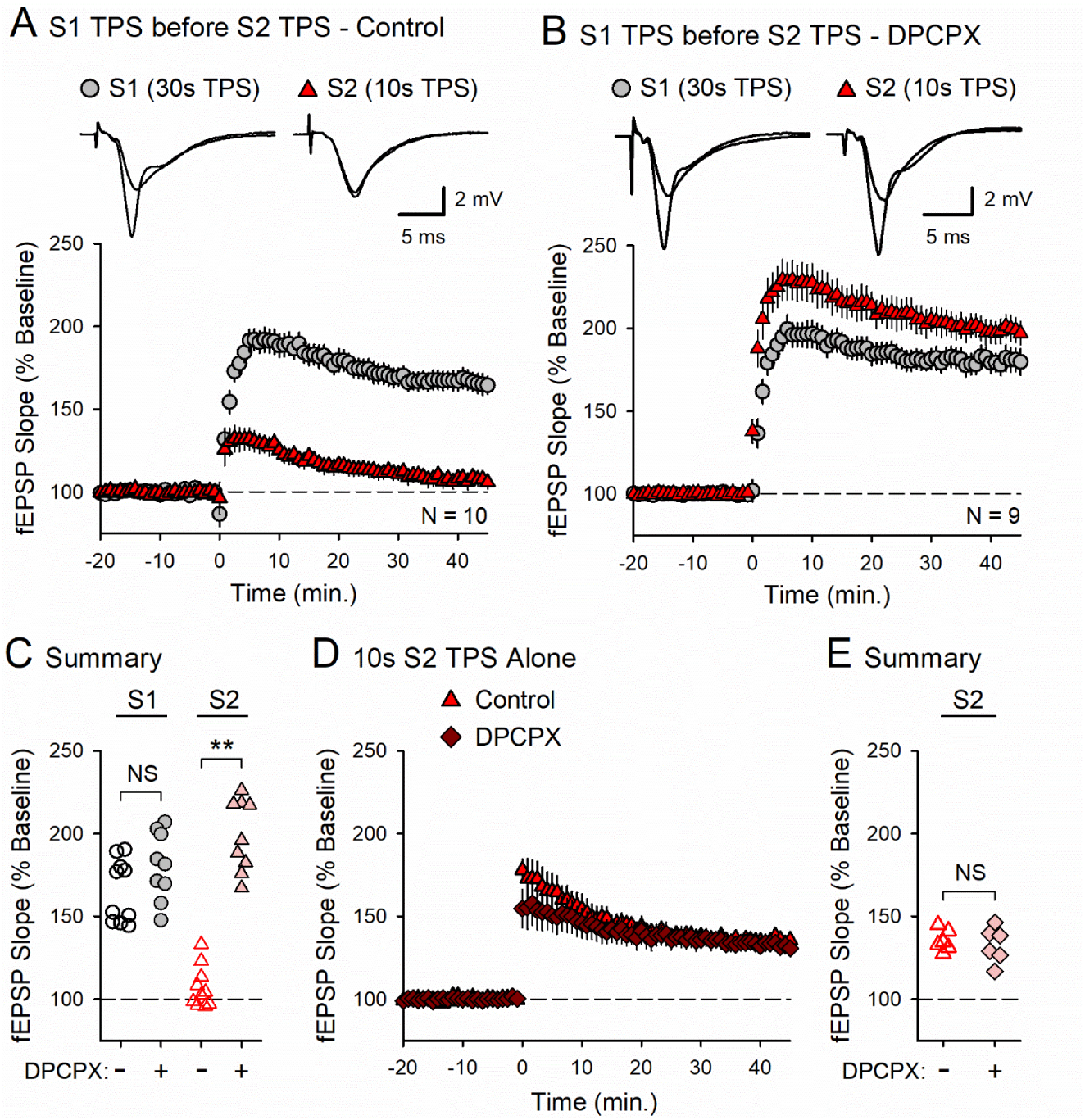
TPS induces an A1 receptor-dependent form of BCM-like metaplasticity. (***A***) In control experiments, 30 seconds of S1 TPS delivered before a 10-seconds-long train of S2 TPS (delivered at time = 0) inhibited BTSP induction at S2 synapses. 45 minutes after TPS S1 fEPSPs were potentiated to 165 ± 6% of baseline and S2 fEPSPs were just 107 ± 6% of baseline. (***B***) DPCPX blocks the heterosynaptic suppression of BTSP induced by S1 TPS. 30 seconds of S1 TPS delivered before 10 seconds of S2 TPS induced a robust potentiation of both S1 and S2 synapses in slices continuously bathed in ACSF containing 400 nM DPCPX. 45 minutes post-TPS S1 synapses potentiated to 180 ± 7% of baseline and S2 synapses potentiated to 199 ± 7% of baseline. Traces in A and B show superimposed fEPSPs recorded during baseline and 45 minutes-post TPS. (***C***) S1 and S2 fEPSP slopes 45 minutes after TPS from all experiments in A and B. A two-way ANOVA revealed a significant effect of synapse, *F*_(1,34)_ = 10.665, *p* = 0.002, a significant effect of DPCPX, *F*_(1,34)_ = 77.399, *p* < 0.001, and a significant interaction, *F*_(1,34)_ = 40.367, *p* < 0.001. DPCPX significantly enhanced BTSP induction at S2 synapses (\*\**p* < 0.001) but had no effect on the induction of BTSP at S1 synapses (NS not significant, *p* = 0.093). (***D***) 10 seconds of S2 TPS was delivered (at time = 0) in the absence (control, n = 6) or presence of 400 nM DPCPX (n = 6). 45 minutes post-TPS fEPSPs were potentiated to 135 ± 3% of baseline in control experiments (n = 6) and 133 ± 4% of baseline in experiments where slices were continuously bathed in ACSF containing DPCPX. (***E***) Summary showing fEPSP slopes 45 minutes post-S2 TPS from all experiments in D (NS not significant, *t*_(10)_ = 0.506, *p* = 0.623).

## Discussion

The hypothesis that burst-dependent forms of plasticity are regulated by a novel, burst-dependent form of BCM plasticity (Payeur et al., 2021) makes two readily testable predictions. First, the induction of burst-dependent forms of plasticity, such as BTSP, at one group of synapses should increase the threshold for activity-dependent increases in synaptic strength at other synapses. Second, unlike standard BCM plasticity, activity-dependent changes in θ_M_ should be triggered by postsynaptic bursts, rather than changes in the frequency of single postsynaptic action potentials. Consistent with the first of these predictions, patterns of TPS that induce robust homosynaptic BTSP also trigger a pronounced, heterosynaptic suppression of BTSP induction. Consistent with the second, the A1 adenosine receptor-mediated heterosynaptic depression that underlies the heterosynaptic suppression of BTSP induction is a CS burst-dependent form of plasticity.

Interestingly, postsynaptic CS bursts not only provide the postsynaptic depolarization needed for NMDA receptor activation and BTSP induction (O’Dell, 2022) but, as shown here, also provide the postsynaptic depolarization needed for activation of L-type Ca^2+^ channels and the induction of heterosynaptic depression. However, while blocking L-type channels with nimodipine inhibits the induction of heterosynaptic depression it does not inhibit EPSP-evoked CS bursting during TPS (Fig. 6A). This is consistent with previous findings indicating that although L-type Ca^2+^ channels are present in both the dendritic spines and dendrites of CA1 pyramidal cells (Tippens et al., 2008; Leitch et al., 2009; Davare et al., 2001), these channels are not required for the generation of dendritic plateau potentials (Takahashi and Magee, 2009; Bittner et al., 2017). Thus, rather than having an important role in generating the dendritic plateau potentials and somatic CS bursts needed for the induction of BTSP, L-type channels may instead serve as “burst-detectors” responsible for the induction of an A1 receptor and burst-dependent BCM-like metaplasticity. Consistent with this notion, nimodipine not only blocks TPS-induced heterosynaptic depression but has also been shown to inhibit the activity-dependent release of adenosine from CA1 pyramidal cells (Yamashiro et al., 2017; Wu et al., 2023).

How might an L-type Ca^2+^ channel-dependent increase in intracellular Ca^2+^ lead to increases in extracellular adenosine during a bout of CS bursting? Although intracellular adenosine levels are regulated by a complex set of metabolic pathways, activity-dependent increases in intracellular adenosine levels in neurons are primarily generated by the hydrolysis of ATP (Garcia-Gil et al., 2021; Yahiro et al., 2025; Wei et al., 2025). Thus, L-type Ca^2+^ channel-mediated increases in intracellular Ca^2+^ and the subsequent hydrolysis of ATP by Ca^2+^-ATPases working to restore basal Ca^2+^ levels may have an important role in elevating intracellular levels of adenosine in response to postsynaptic CS bursts. Subsequent release of adenosine into the extracellular space via passive transport mediated by equilibrative nucleoside transporters (Lovatt et al., 2012; Yamashiro et al., 2017; Wu et al., 2023) could then produce the A1 receptor activation responsible for CS burst-dependent heterosynaptic depression and the BCM-like metaplastic suppression of BTSP. A number of studies have found, however, that A1 receptor-dependent forms of synaptic depression in the hippocampal CA1 region may involve ATP release from astrocytes, followed by the enzymatic conversion of extracellular ATP to adenosine by ecto-nucleotidases (Pascual et al., 2005; Serrano et al., 2006; Andersson et al., 2008; Wall and Dale, 2013). Microglia may also have an important role in adenosine signaling in the CNS (Badimon et al., 2020). Thus, although recent studies using genetically encoded adenosine sensors indicate that neurons are the primary source for activity-dependent increases in extracellular adenosine (Wu et al., 2023; Wei et al., 2025, also see Lovatt et al., 2012), the potential role of glial cells in burst-dependent forms of metaplasticity is an interesting question to investigate in future studies.

Although the burst-dependent release of adenosine from neurons and activation of presynaptic A1 receptors is a retrograde form of synaptic transmission, A1 receptors are also found in the dendritic spines and dendrites of CA1 pyramidal cells (Ochiishi et al., 1999; Rebola et al., 2003). The dendritic spines and dendrites of CA1 pyramidal cells also contain GIRK-type K^+^ channels (Drake et al., 1997; Kulik et al, 2006) that are activated by G protein-coupled receptors, including A1 receptors (Trussell and Jackson, 1987; Luscher et al., 1997). Together, these findings suggest that the postsynaptic hyperpolarization produced by adenosine-activated GIRK channels can also contribute to the heterosynaptic inhibition of BTSP induction by opposing the membrane depolarization needed to relieve the voltage-dependent Mg^2+^ block of NMDA receptor ion channels. Thus, rather than acting as a retrograde messenger, adenosine likely instead acts as a paracrine factor that suppresses NMDA receptor activation and the induction of BTSP by activating both pre- and postsynaptic A1 receptors.

In addition to using CS bursts to dynamically regulate θ_M_, the burst-dependent metaplasticity induced by TPS differs from conventional BCM plasticity in other important ways. For example, in standard BCM plasticity the induction of LTP at one group of synapses is associated with a heterosynaptic shift in θ_M_ that not only suppresses LTP induction but also enhances the induction of LTD. However, although 30 seconds of TPS induced a heterosynaptic suppression of BTSP induction, it did not facilitate the induction of LTD (Fig. 2D, Supplemental Fig. S3). Because standard protocols for the induction of NMDA receptor-dependent LTD typically use 900 pulses of low-frequency presynaptic fiber stimulation (Dudek and Bear, 1992), the comparatively brief, 100-pulse trains of stimulation used in the experiments shown in figure 2 may have simply been below threshold for LTD induction, even after the heterosynaptic shift in θ_M_ induced by 30 seconds of TPS. Alternatively, the apparent absence of facilitated LTD induction following 30 seconds of TPS may reflect a fundamental difference between burst-dependent and conventional forms of BCM metaplasticity. Indeed, unlike Hebbian plasticity, BTSP is a weight-dependent form of plasticity where strong synapses depress (and weak synapses potentiate) in response to appropriately timed dendritic plateau potentials (Milstein et al. 2021; Magee, 2026). It will thus be interesting to explore how burst-dependent BCM-like metaplasticity regulates the plasticity of potentiated synapses in future experiments.

The brief duration of CS burst-dependent metaplasticity (Fig. 3) is also strikingly different from the persistent BCM-like suppression of Hebbian LTP produced by high-frequency stimulation protocols, which lasts for at least 30–60 minutes in-vitro (Wang and Wagner, 1999; Le Ray et al., 2004; Hulme et al., 2012; Hegemann and Abraham, 2021) and several days in-vivo (Abraham et al., 2001). Importantly, activity-dependent shifts in θ_M_ in standard BCM plasticity are induced when a moving average of ongoing spike rates deviates from a homeostatic setpoint or target rate. Standard BCM plasticity is thus a form of homeostatic plasticity where plasticity thresholds are modified to produce changes in synaptic weights that achieve firing rate homeostasis (Keck et al., 2017; Zenke and Gerstner, 2017). In contrast, the relatively short-lived shift in θ_M_ induced by CS bursts suggests that burst-dependent BCM-like plasticity acts as a transient, negative-feedback form of plasticity that is primarily involved in generating competitive synaptic interactions specifically during bouts of CS bursting. Thus, rather than acting as a long-term form of homeostatic plasticity, the transient heterosynaptic suppression of BTSP induced by CS bursts likely instead provides a mechanism that promotes sparse encoding of information by limiting the number of synapses undergoing BTSP during bouts of theta-frequency CS bursting. Notably, theta-frequency CS bursting is prominent in CA1 pyramidal cells during spatial learning (Otto et al., 1991; Tanaka et al., 2018) and the induction of BTSP by dendritic plateau potentials and somatic CS bursts underlies the for the initial formation of hippocampal place cells during spatial learning (Bittner et al., 2017; Milstein et al., 2021; Magee, 2026). Theta-frequency CS bursting is, however, also prominent during place field firing in CA1 pyramidal cells with established place fields (Epsztein et al., 2011; Lowet et al., 2023). Thus, CS burst-dependent BCM-like metaplasticity may be re-engaged in familiar environments to help maintain sparse encoding of spatial information during place-specific firing in established place cells.

## Materials and Methods

### Slice Preparation

Dorsal hippocampal slices were obtained from 8-14 weeks old, male and female C57Bl/6 mice (#027 Charles River Laboratories) using techniques approved by the Institutional Animal Care and Use Committee at the University of California, Los Angeles. Following isoflurane anesthesia and subsequent cervical dislocation, the brain was removed and submerged in a cold (∼4°C), oxygenated (95% O_2_/5% CO_2_) ACSF containing 124 mM NaCl, 4 mM KCl, 25 mM NaHCO_3_, 1 mM NaH_2_PO_4_, 2 mM CaCl_2_, 1.2 mM MgSO_4_, and 10 mM glucose. After allowing the brain to cool for 5 minutes in cold ACSF, hippocampi were dissected from the brain, and a manual tissue slicer (Stoelting Co.) was used to prepare 400-µm-thick slices. A micro-knife (Fine Science Tools) was used to remove the CA3 region and slices were transferred into interface-type chambers continuously perfused with ACSF (2-3 mL/min at 30°C) and allowed to recover for 2-6 hours before recordings began.

### Slice Electrophysiology

Slices were maintained at 30°C in an interface-type recording chamber perfused with ACSF. Two bipolar stimulating electrodes (S1 and S2) manufactured from twisted strands of Formvar-coated, nickel-chromium wire (A-M Systems) were placed in stratum radiatum (one near the CA3 region and the other near the subiculum) to activate independent groups of Schaffer collateral/commissural fiber synapses onto CA1 pyramidal cells. fEPSPs were recorded using a glass microelectrode (filled with ACSF, resistance ≍ 10 MΩ) placed in stratum radiatum between the stimulating electrodes. fEPSPs were evoked using 0.02 msec duration voltage pulses, low-pass filtered at 2 kHz and digitized at 10 kHz using a Multi-Clamp 700B amplifier (Molecular Devices). At the start of each experiment the maximal amplitude of fEPSPs evoked by each stimulating electrode was determined and the stimulation intensity was adjusted to evoke fEPSPs that were approximately 50% of the maximal amplitude. Standard techniques described elsewhere (O’Dell, 2022) were used to confirm activation of independent groups of synapses and alternating pulses of S1 and S2 presynaptic fiber stimulation were then delivered every 25 seconds. In experiments investigating heterosynaptic depression S2 synapses were activated once every 30 seconds before and after different trains of stimulation delivered to S1 synapses. All chemicals were obtained from Millipore-Sigma except for TTX citrate (Alomone Labs) and DAMGO (Hello Bio). Stock solutions of TTX (1 mM), and DAMGO (1 mM) were prepared in dH_2_O and stored at −20°C. Stock solutions of 8-cyclopentyl-1,3-dipropylxanthine (DPCPX, 2 mM) and nimodipine (10 mM) were prepared in DMSO and stored at −20°C.

BTSP was induced using theta-pulse stimulation (TPS) protocols (single pulses of 5 Hz presynaptic fiber stimulation). Note that unlike more commonly used theta-burst stimulation protocols, where Hebbian LTP is induced by brief bursts of 100 Hz stimulation delivered at 5 Hz (Larson and Munkácsy, 2015), TPS uses single stimulation pulses. Thus, unlike standard high-frequency stimulation protocols, TPS protocols induce BTSP by evoking CS bursts in CA1 pyramidal cells (Thomas et al., 1998; O’Dell, 2022) and thus generate, in-vitro, the prominent theta-frequency CS bursting that occurs in CA1 pyramidal cells in-vivo during memory encoding (Otto et al., 1991; Tanaka et al., 2018).

### Statistical Analysis

fEPSPs were analyzed using pClamp10.7 software (Molecular Devices) and SigmaPlot 12.5 (Grafiti) was used for statistical tests. Average slopes of fEPSPs (normalized to baseline) recorded 40-45 minutes post-TPS were used for statistical comparisons. EPSP-evoked CS bursting during TPS was quantified by counting the number of negative-going population spikes (PSs) elicited by each EPSP during TPS and the probability of EPSP-evoked CS bursting (*P*_CSB_) was determined from the number of EPSPs eliciting 2 or more PSs relative to the total number of EPSPs evoked during TPS. The slope of the first S2 fEPSPs (normalized to baseline) elicited after stimulation trains delivered to S1 synapses was used to measure heterosynaptic depression. Student’s t-tests were used to evaluate statistical significance between two groups. Multiple comparisons were done using one-way or two-way ANOVAs with Student-Newman-Keuls *post hoc* comparisons. Pearson Product Moment Correlation tests were used to evaluate correlations. No obvious sex differences were found and results from male and female mice were combined. Unless noted otherwise, results are reported as mean ± SEM.

## Acknowledgements

This work was supported by funds provided by the UCLA Academic Senate.

---

**Supplemental figure S1:**
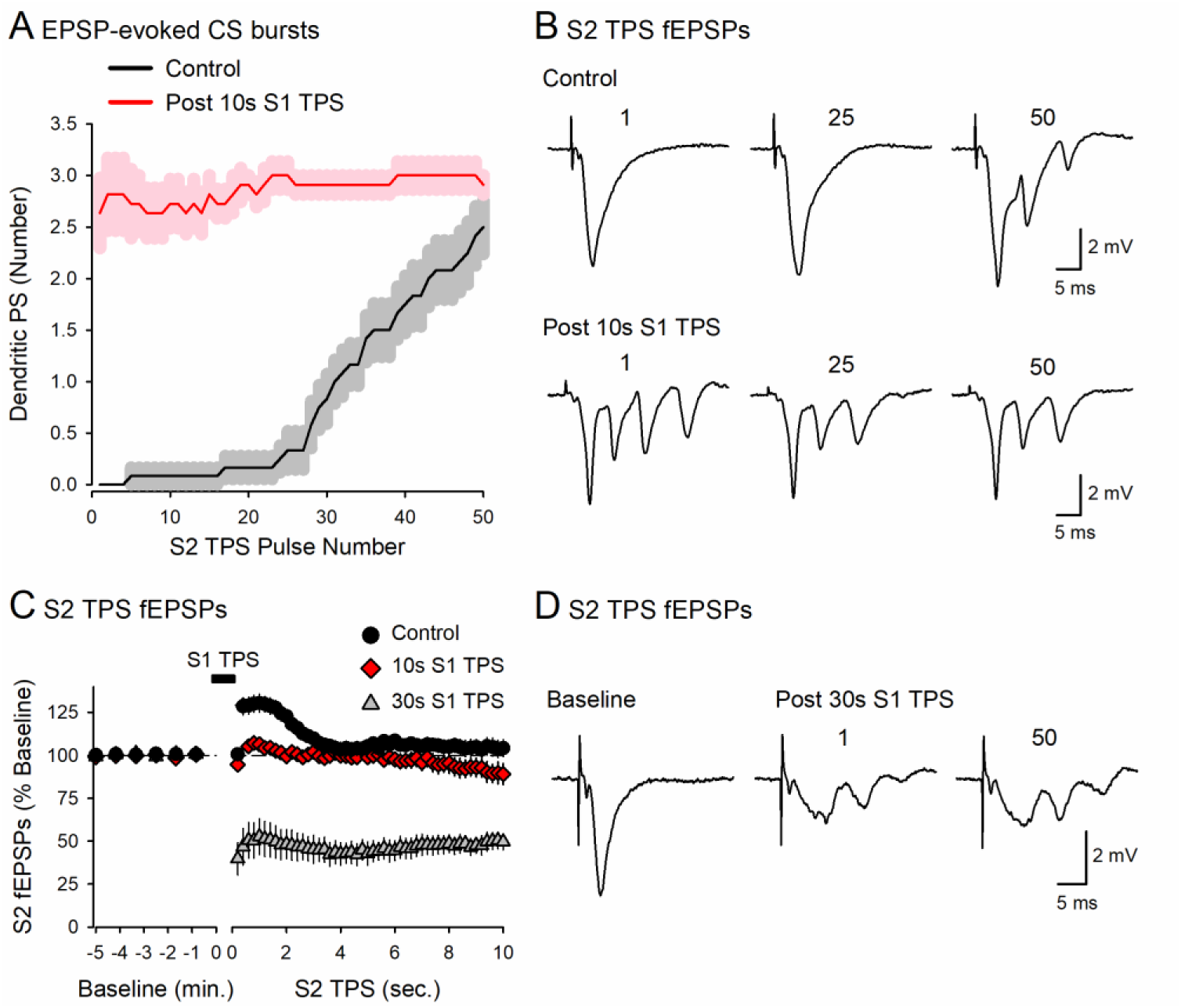
TPS induces multiple forms of heterosynaptic plasticity. (***A***) Plot shows the number of dendritic population spikes (PS) elicited by EPSPs during 10 seconds of TPS delivered to S2 synapses. Shading represents ± SEM. In control experiments, where S2 TPS was delivered without prior S1 stimulation, EPSP-evoked CS bursting (defined as fEPSPs that elicit 2 or more negative-going dendritic PS) developed in a highly activity-dependent manner and only began after ∼7-8 seconds of TPS (n = 12). In contrast, 10 seconds of TPS delivered to S1 synapses before S2 TPS induced a robust heterosynaptic facilitation EPSP-evoked CS bursting during S2 TPS (n = 11). (***B***) Traces show fEPSPs evoked by the 1^st^, 25^th^, and 50^th^ pulse of S2 TPS in a control experiment and in an experiment where 10 seconds of S1 TPS was delivered before S2 TPS. (***C***) S2 fEPSPs recorded during S2 TPS in control experiments and in experiments where either 10 or 30 seconds of S1 TPS were delivered before S2 TPS (n’s = 12, 11, and 9, respectively). Note the strong heterosynaptic depression of S2 fEPSPs induced by 30 seconds of S1 TPS. (***D***) Traces show S2 fEPSPs recorded during baseline and elicited by the indicated pulse numbers during a train of S2 TPS that was delivered after 30 seconds of S1 TPS. Results are from the same experiments shown in figure 1A-C.

**Supplemental figure S2:**
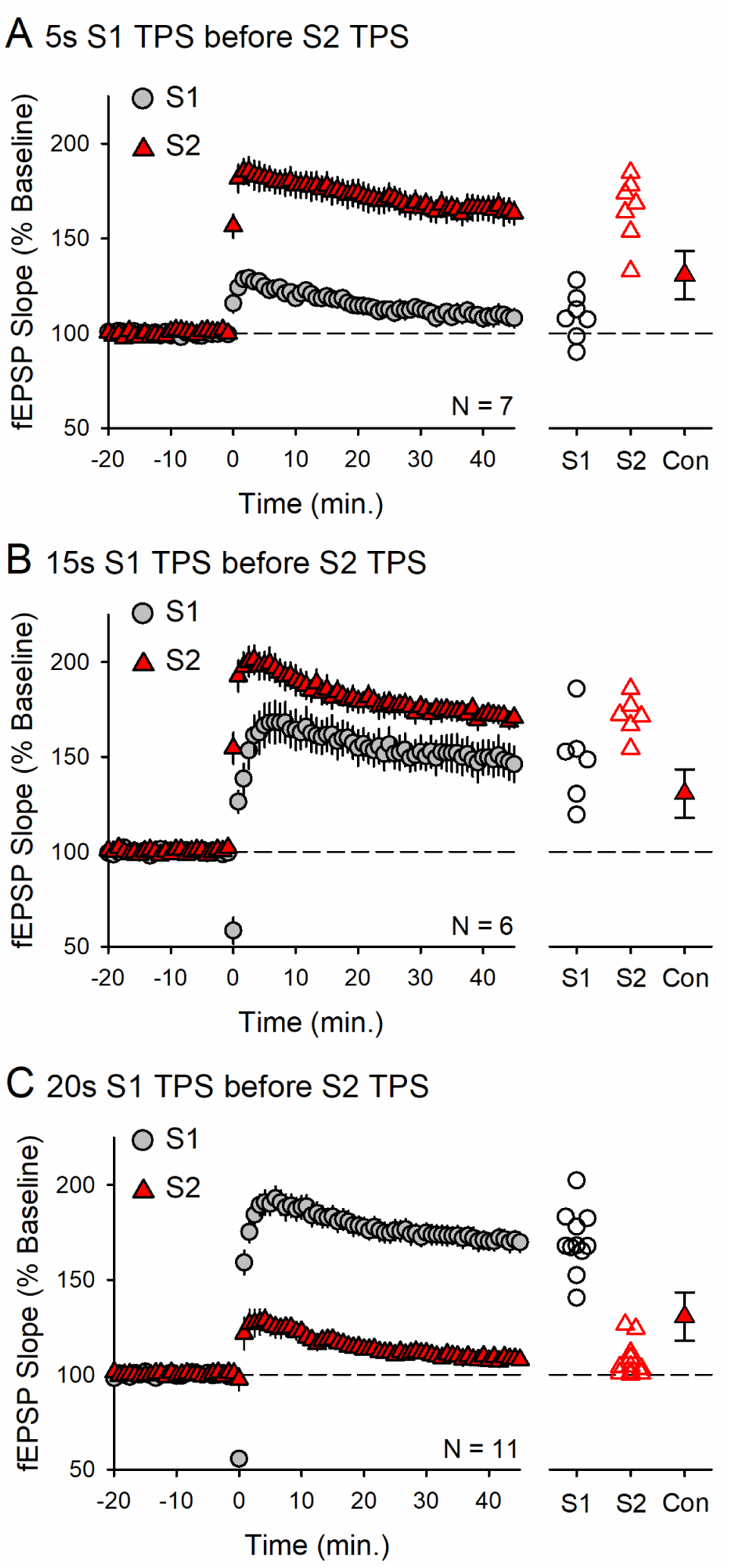
Additional results for figure 1D. (***A***) Five seconds of S1 TPS delivered before 10 seconds of S2 TPS (at time = 0) had little lasting effect on synaptic strength at S1 synapses (45 minutes post-TPS S1 fEPSPs were 109 ± 5% of baseline) but facilitated BTSP induction at S2 synapses (45 minutes post-TPS S2 synapses were potentiated to 165 ± 7% of baseline). (***B***) 15 seconds of S1 TPS delivered before 10 seconds of S2 TPS (at time = 0) facilitates BTSP induction at S2 synapses (45 minutes post-TPS S1 fEPSPs were potentiated to 149 ± 9% of baseline and S2 synapses potentiated to 171 ± 4% of baseline). (***C***) Twenty seconds of S1 TPS delivered before 10 seconds of S1 TPS induces BTSP at S1 synapses (S1 fEPSPs potentiated to 170 ± 5% of baseline) and suppresses BTSP induction at S2 synapses (45 minutes post-TPS S2 fEPSPs were 107 ± 3% of baseline). Scatter plots in A-C show S1 and S2 fEPSP slopes recorded 45 minutes post-TPS from all experiments. For comparison, the filled triangles show the average (± standard deviation) potentiation induced by 10 seconds of TPS delivered to S2 synapses alone in the control (Con) experiments shown in figure 1A.

**Supplemental figure S3:**
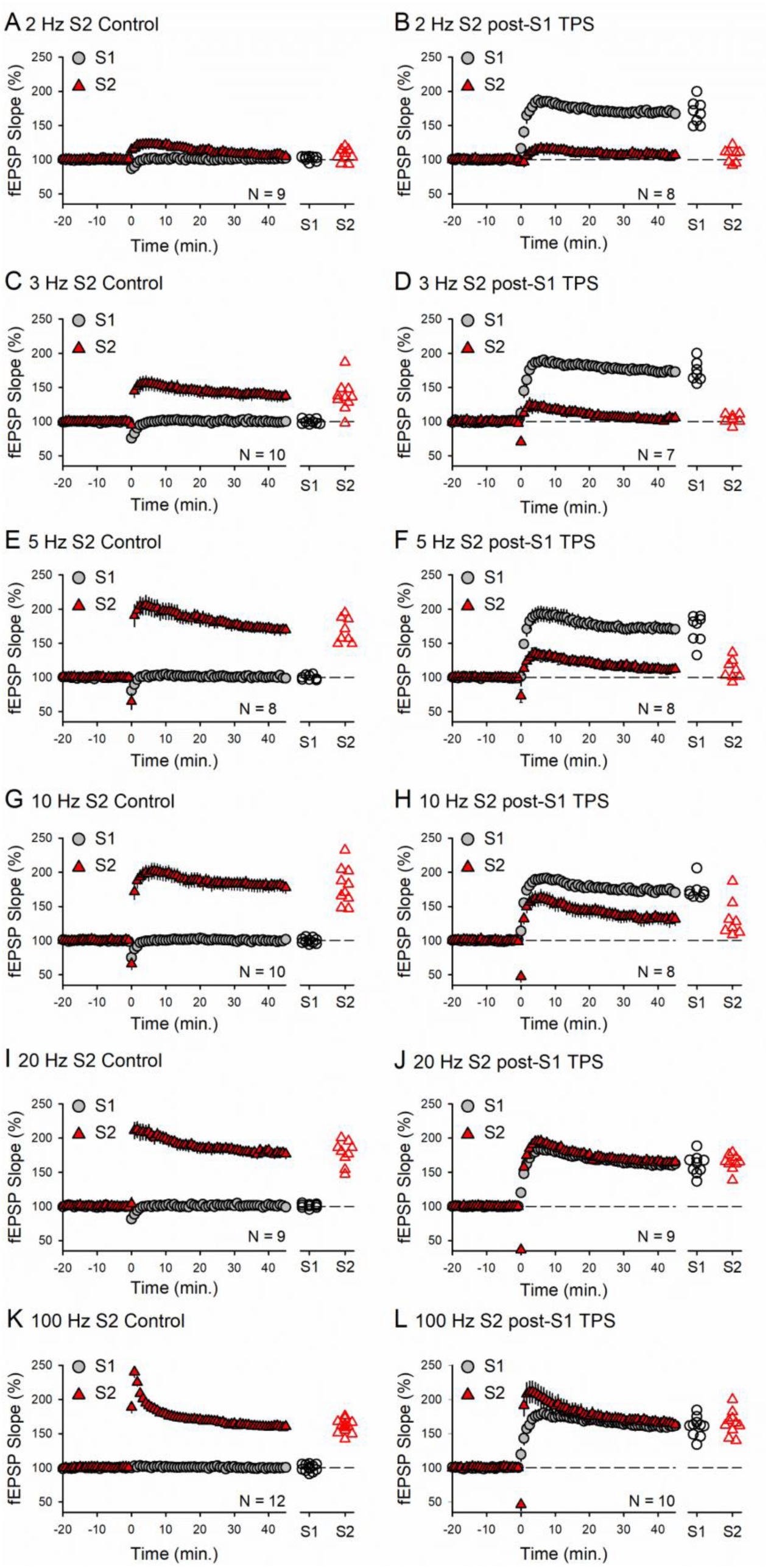
Additional results for figure 2D. 100 pulses of S2 stimulation were delivered (at time = 0) at the indicated frequencies either alone (left column) or after a 30 seconds-long train of TPS delivered to S1 synapses (right column). Scatter plots show average S1 and S2 fEPSP slopes recorded 45 minutes after stimulation protocols from all experiments.

**Supplemental figure S4:**
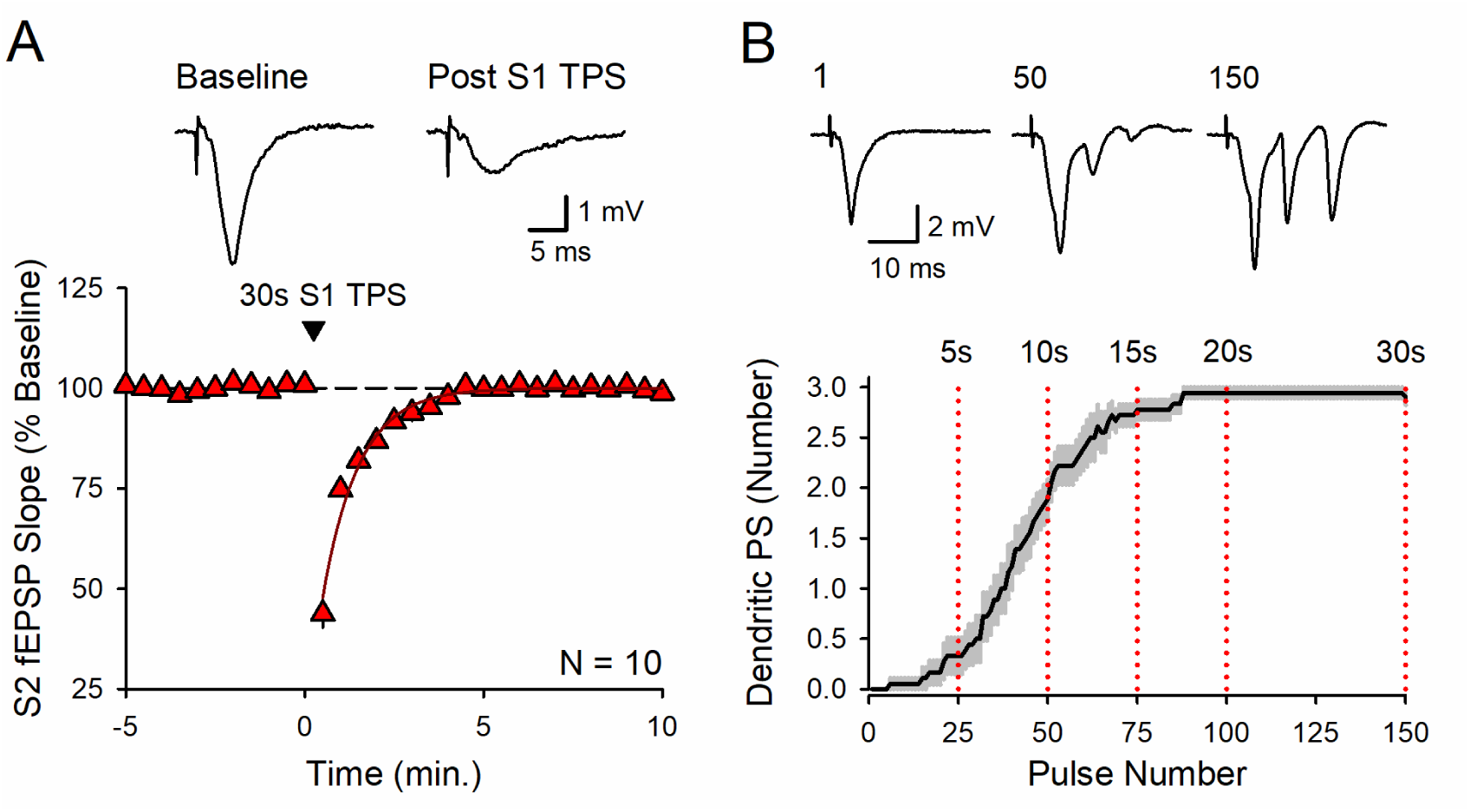
Heterosynaptic depression and EPSP-evoked CS bursting induced by 30 seconds of S1 TPS. (***A***) TPS induces short-term heterosynaptic depression. S2 fEPSPs were evoked at 0.033 Hz before and after 30 seconds of S1 TPS (delivered at time = 0). Traces show S2 fEPSPs recorded during baseline and the first S2 fEPSP evoked post-S1 TPS. Following S1 TPS, S2 fEPSPs were initially depressed to 44 ± 3% of baseline. Line shows a single exponential fit to the recovery of S2 fEPSPs following S1 TPS (τ = 60.5 ± 4s). (***B***) Number of dendritic populations spikes (PS) evoked by EPSPs during 30 seconds of S1 TPS. Shading represents ± SEM (n = 10). Traces show examples of fEPSPs elicited by the 1^st^, 50^th^, and 150^th^ stimulation pulses during TPS. Note the activity-dependent development of negative-going PSs.

**Supplemental figure S5:**
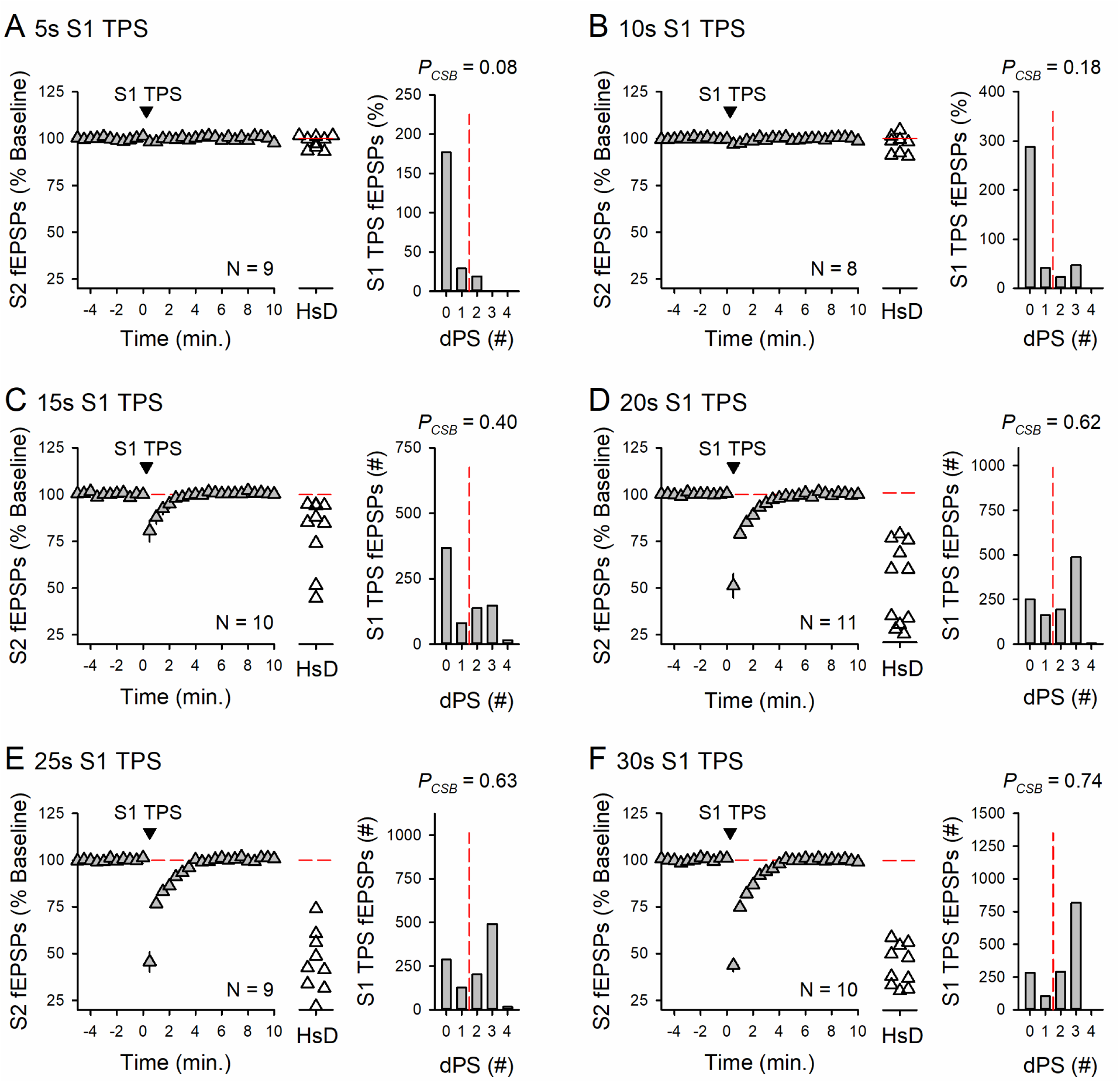
Additional results for Figure 4B. Plots show the effects of the indicated durations of S1 TPS (delivered at time = 0) on synaptic transmission at S2 synapses. Scatter plots show the slopes of the first S2 fEPSPs elicited post-S1 TPS used to measure heterosynaptic depression (HsD) from all experiments. Histograms show the number S1 EPSPs eliciting 0-4 dendritic population spikes (dPS) during S1 TPS in all experiments. Note how the probability of EPSP-evoked CS bursting (*P_CSB_*) during S1 TPS and the magnitude of heterosynaptic depression at S2 synapses grow as the duration of S1 TPS increases.

**Supplemental figure 6:**
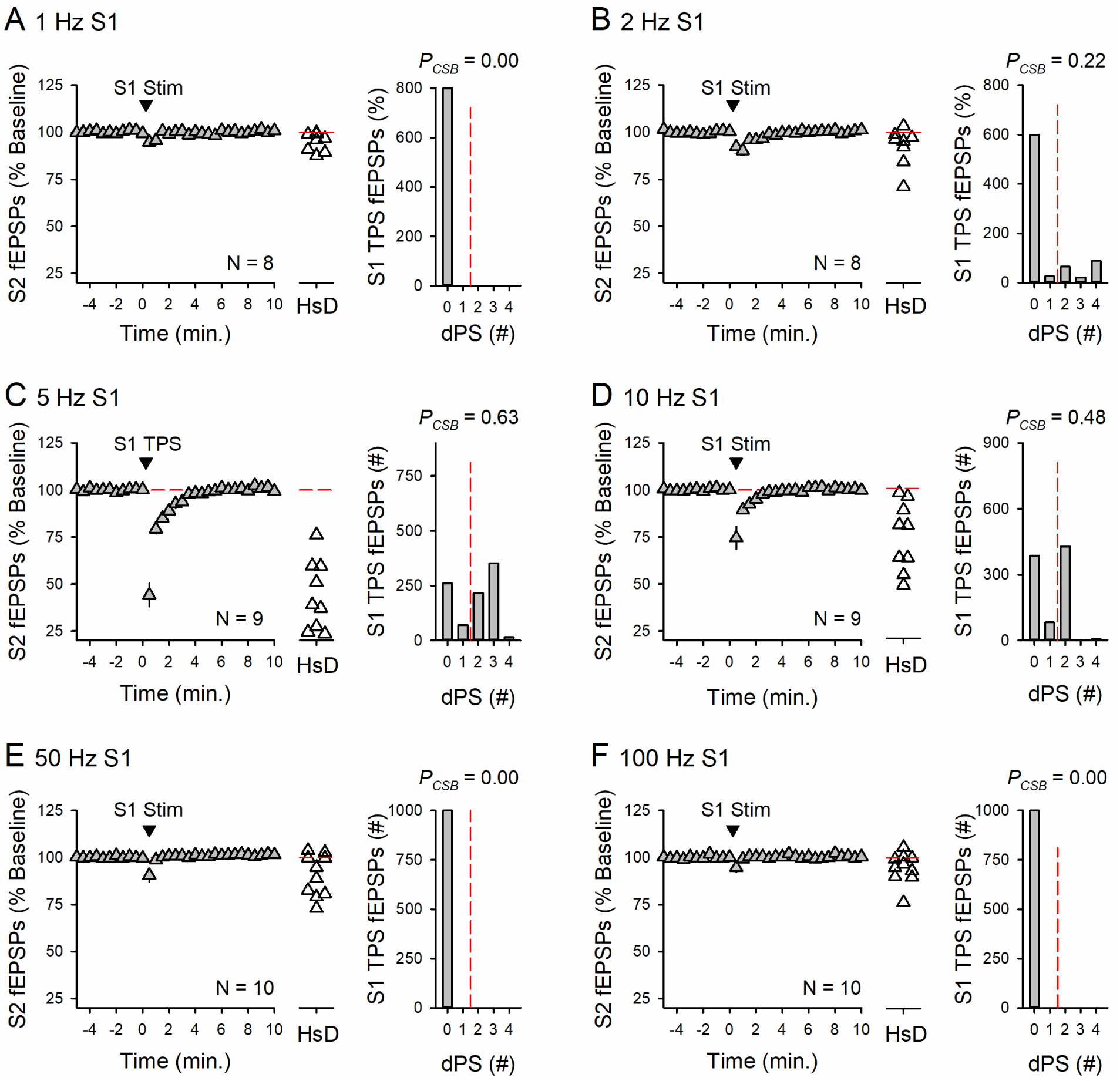
Additional results for Figure 4D. Plots show the effects of 100 pulse trains of 1-100 Hz S1 stimulation (delivered at time = 0) on synaptic transmission at S2 synapses. Scatter plots show the slopes of the first S2 fEPSPs elicited post-S1 stimulation trains used to measure heterosynaptic depression (HsD) from all experiments. Histograms show the number S1 EPSPs eliciting 0-4 dendritic population spikes (dPS) during S1 stimulation in all experiments. *P_CSB_* is the probability of EPSP-evoked CS bursting during S1 stimulation.

**Supplemental figure 7:**
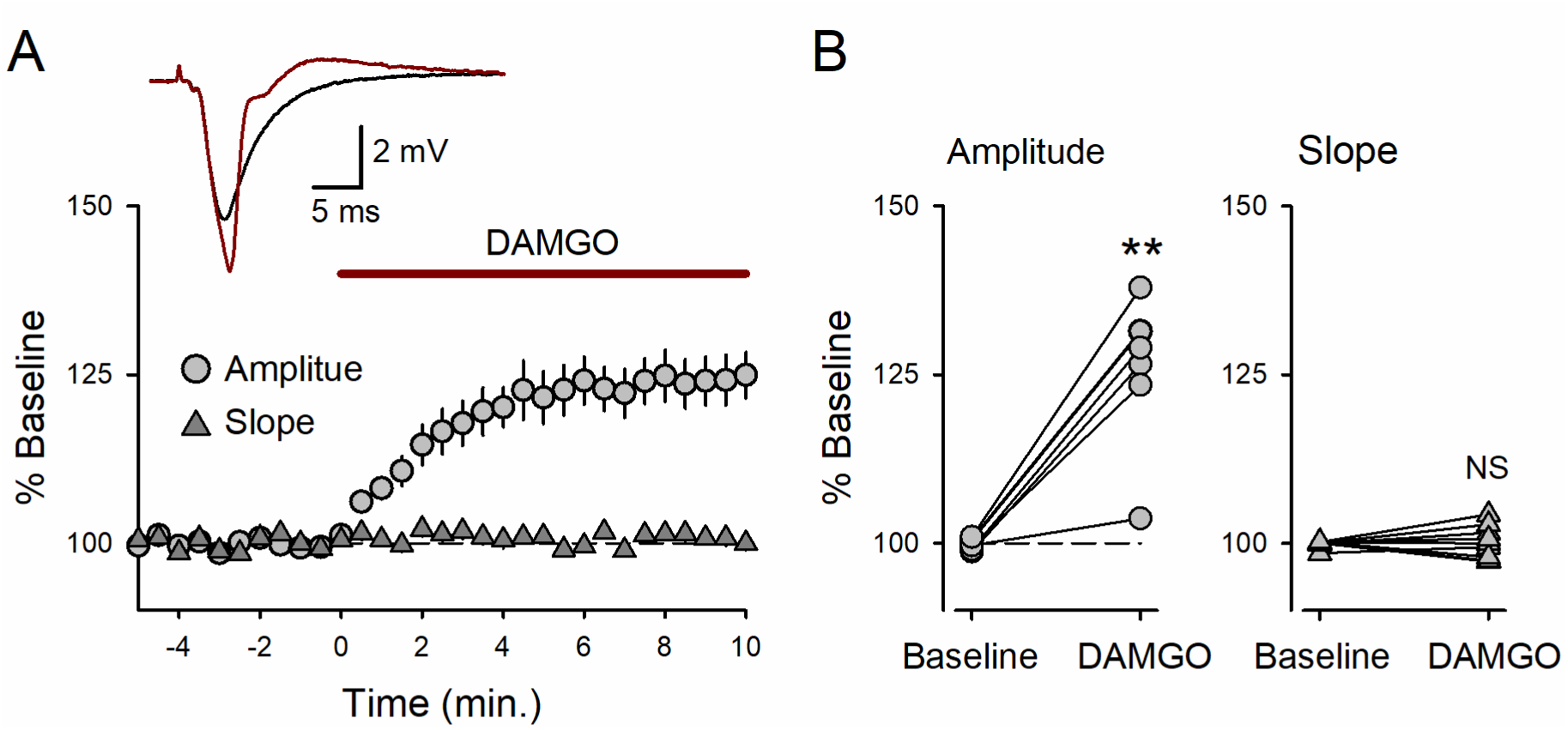
(***A***) Bath application of the µOR receptor agonist DAMGO increases the amplitude of fEPSPs but has no effect on fEPSP slopes. Traces show superimposed fEPSPs recorded during baseline and at the end of a 10-minute application of 1 µM DAMGO (indicated by the bar). (***B***) Results from all experiments (n = 8) showing the effects of µOR activation on fEPSP amplitudes (left) and fEPSP slopes (right). DAMGO significantly increased fEPSP amplitudes (*t*_(7)_ = 6.954, \*\**p* = 0.00022, paired t-test comparison to baseline) but had no effect on fEPSP slopes (*t*_(7)_ = 0.746, NS not significant, *p* = 0.48). Average fEPSP amplitudes and slopes during the last minute of baseline and last minute of DAMGO application were used for statistical comparisons.

